# Intronic elements associated with insomnia and restless legs syndrome exhibit cell type-specific epigenetic features contributing to *MEIS1* regulation

**DOI:** 10.1101/2021.09.03.458823

**Authors:** Daniel D. Lam, Ana Antic Nikolic, Chen Zhao, Nazanin Mirza-Schreiber, Wojciech Krężel, Konrad Oexle, Juliane Winkelmann

## Abstract

A highly evolutionarily conserved *MEIS1* intronic region is strongly associated with restless legs syndrome (RLS) and insomnia. To understand its regulatory function, we dissected the region by analyzing chromatin accessibility, enhancer-promoter contacts, DNA methylation, and eQTLs in different human neural cell types and tissues. We observed specific activity with respect to cell type and developmental maturation, indicating a prominent role for distinct highly conserved intronic elements in forebrain inhibitory neuron differentiation. Two elements were hypomethylated in neural cells with higher *MEIS1* expression, suggesting a role of enhancer demethylation in gene regulation. *MEIS1* eQTLs showed a striking modular chromosomal distribution, with forebrain eQTLs clustering in intron 8/9. CRISPR interference targeting of individual elements in this region attenuated *MEIS1* expression, revealing a complex regulatory interplay of distinct elements. In summary, we found that *MEIS1* regulation is organized in a modular pattern. Disease-associated intronic regulatory elements control *MEIS1* expression with cell type and maturation stage specificity, particularly in the inhibitory neuron lineage. The precise spatiotemporal activity of these elements likely contributes to the pathogenesis of insomnia and RLS.

## INTRODUCTION

A series of genome-wide association studies (GWAS) consistently identify strong association signals for restless legs syndrome (RLS), a common neurological disorder, in introns 7-9 of the *MEIS1* locus (Schormair et al. 2017; Winkelmann et al. 2011, 2007). A recent series of GWAS for insomnia, sleep, and circadian traits also identify strong association signals in this same region, in part due to phenotypic overlap with RLS (Lane et al. 2019; Jones et al. 2019; Lane et al. 2017; Jansen et al. 2019; Hammerschlag et al. 2017).

*MEIS1* encodes a homeodomain-containing transcription factor of the three amino acid loop extension (TALE) superclass (Schulte and Geerts 2019). MEIS proteins are broadly expressed in embryonic development and regulate diverse developmental processes, including limb and vascular patterning, eye and neural development, hematopoiesis, and cardiogenesis (Schulte and Geerts 2019). The scope of these diverse roles in disparate tissues necessitates precise spatiotemporal regulation of *MEIS1* expression. Such precision is typically conferred by arrayed non-coding regulatory elements (Pennacchio et al. 2006; Nord et al. 2013).

Indeed, the *MEIS1* locus harbours an array of highly evolutionarily conserved non-coding elements with presumed regulatory function. Here, we interrogated tissue and cell type-specific chromatin accessibility, promoter-enhancer contacts, and DNA methylation as readouts of functional activity using cultured human cells. We show that several elements in the disease-associated intronic region are active in forebrain neural development, particularly in the inhibitory neuron lineage, with activity peaking during the late neurogenic period. CRISPR interference experiments revealed distributed regulation of *MEIS1* transcription by multiple elements. Thus, intronic elements tightly control *MEIS1* expression in forebrain development, particularly in the inhibitory neuron lineage, and variants within them can impact forebrain development and function *via* transcriptional regulation.

## RESULTS

### Human cell type-specific accessibility of *MEIS1* intronic elements

The accessibility of chromatin is considered to reflect its regulatory capacity (Klemm et al. 2019). Leveraging a recently published atlas of human fetal chromatin accessibility (Domcke et al. 2020), we assessed the *MEIS1* locus in 12 annotated cell types of the cerebellum and cerebrum (telencephalon or forebrain). We observed pronounced cell type specificity of chromatin accessibility, with greatest accessibility of several elements, including those previously designated 602 and 617 (Spieler et al. 2014), in inhibitory neurons of the cerebrum (Fig. 1).

**Figure 1.**
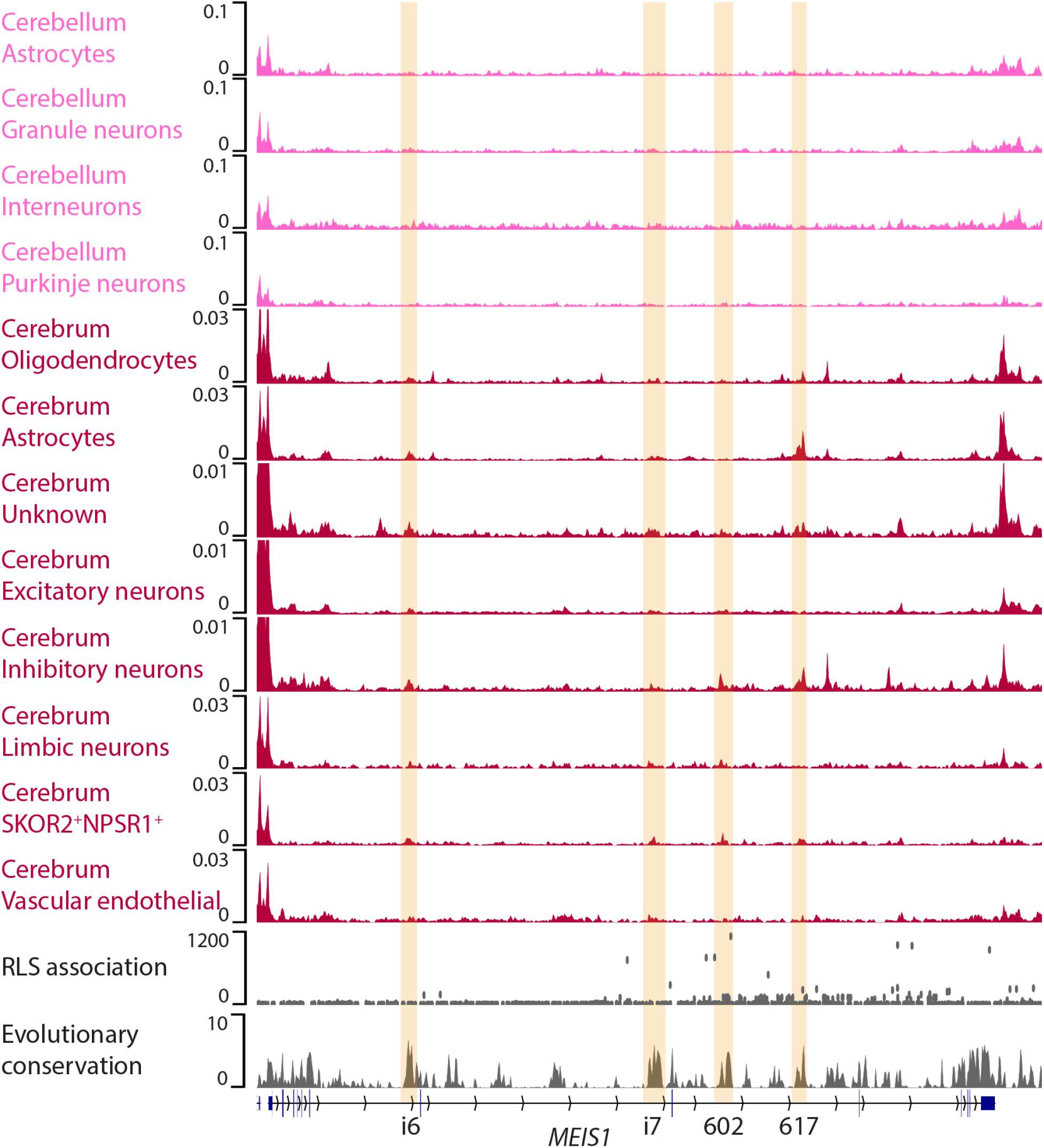
Chromatin accessibility in human brain cell types. Aggregated single cell ATAC-seq read density in 12 fetal human brain cell types. Data from (Domcke et al. 2020). RLS association is from unpublished metaGWAS, scale is -log_10_P. Evolutionary conservation is phyloP 100-way (Pollard et al. 2010). Elements in intron 6 (i6), 7 (i7), and 8 (602 and 617) are highlighted.

Having identified the greater accessibility of intronic *MEIS1* elements in forebrain inhibitory neurons, we sought to further characterize the regulatory landscape of the *MEIS1* locus by implementing *in vitro* differentiation of human induced pluripotent stem cells towards forebrain inhibitory neurons. Forebrain inhibitory neurons are generated in the ganglionic eminences (GEs), transient fetal structures with a well-described architecture (Marín and Rubenstein 2001). GEs are the source of striatal (caudate/putamen) cells, which migrate radially from the ventricular zone towards the striatum as they mature and differentiate, as well as cortical interneurons, which migrate radially and then tangentially towards the cortex (Marín and Rubenstein 2001). In this differentiation paradigm, adapted from (Close et al. 2017), pluripotent stem cells first undergo neural induction towards a forebrain identity, delineated by *FOXG1* and *PAX6* expression (Stoykova et al. 2000; Martynoga et al. 2005), then ventralization towards a GE identity, characterized by *NKX2-1* and *GSX2* expression (Sussel et al. 1999; Corbin et al. 2000) (Fig. 2).

**Figure 2.**
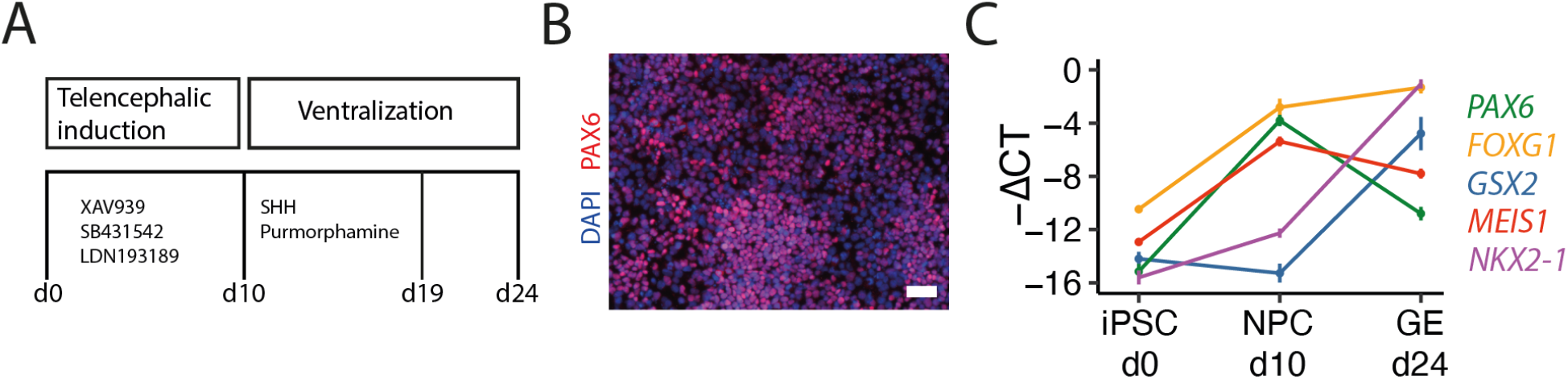
*In vitro* ganglionic eminence differentiation. **A**,*In vitro* ganglionic eminence differentiation scheme. **B**, Successful *in vitro* neural induction shown by PAX6 expression (red). Scale bar, 50 µm. **C**, Expression of key marker genes through differentiation. Values are -(C_T_ of marker gene - C_T_ of *GAPDH* internal reference) and displayed as mean ± SD of 5-6 replicates per stage. C_T_, PCR cycles to threshold.

### Developmental dynamics of chromatin accessibility in the human *MEIS1* locus

To characterize the developmental dynamics of *MEIS1* regulation, we assayed chromatin accessibility using ATAC-seq at three stages of *in vitro* differentiation in human cells: induced pluripotent stem cells (iPSC), neural progenitor cells (NPC), and ganglionic eminence-like cells (GE). We observed changes in accessibility at distinct elements during differentiation (Fig. 3), such as a progressive increase in accessibility of the 617 element and a transient increase in accessibility in an intron 7 element (i7). We also performed ATAC-seq on three commercially available neural cell lines, neural stem cells (NSCs), inhibitory neurons (γ-aminobutyric acid [GABA] neurons), and excitatory glutamatergic neurons. Here too we observed cell type-specific accessibility patterns (Fig. 3). Of note, the 602 element (Spieler et al. 2014), carrying the lead single nucleotide polymorphism (SNP) for RLS association, was only accessible in GABA neurons (Fig. 3). Concordantly, the 602 element was accessible in neural tissue enriched in GABA neurons (fetal lateral ganglionic eminence and putamen, Fig. 3), but not in non-neural cells and tissues where MEIS1 nevertheless has an important function (Fig. 3). An intron 6 element (i6) was also selectively accessible in GABA neurons and NSCs (Fig. 3). Taken together, these results demonstrate selective accessibility of intronic elements in the *MEIS1* locus across different tissues, cell types, and developmental stages.

**Figure 3.**
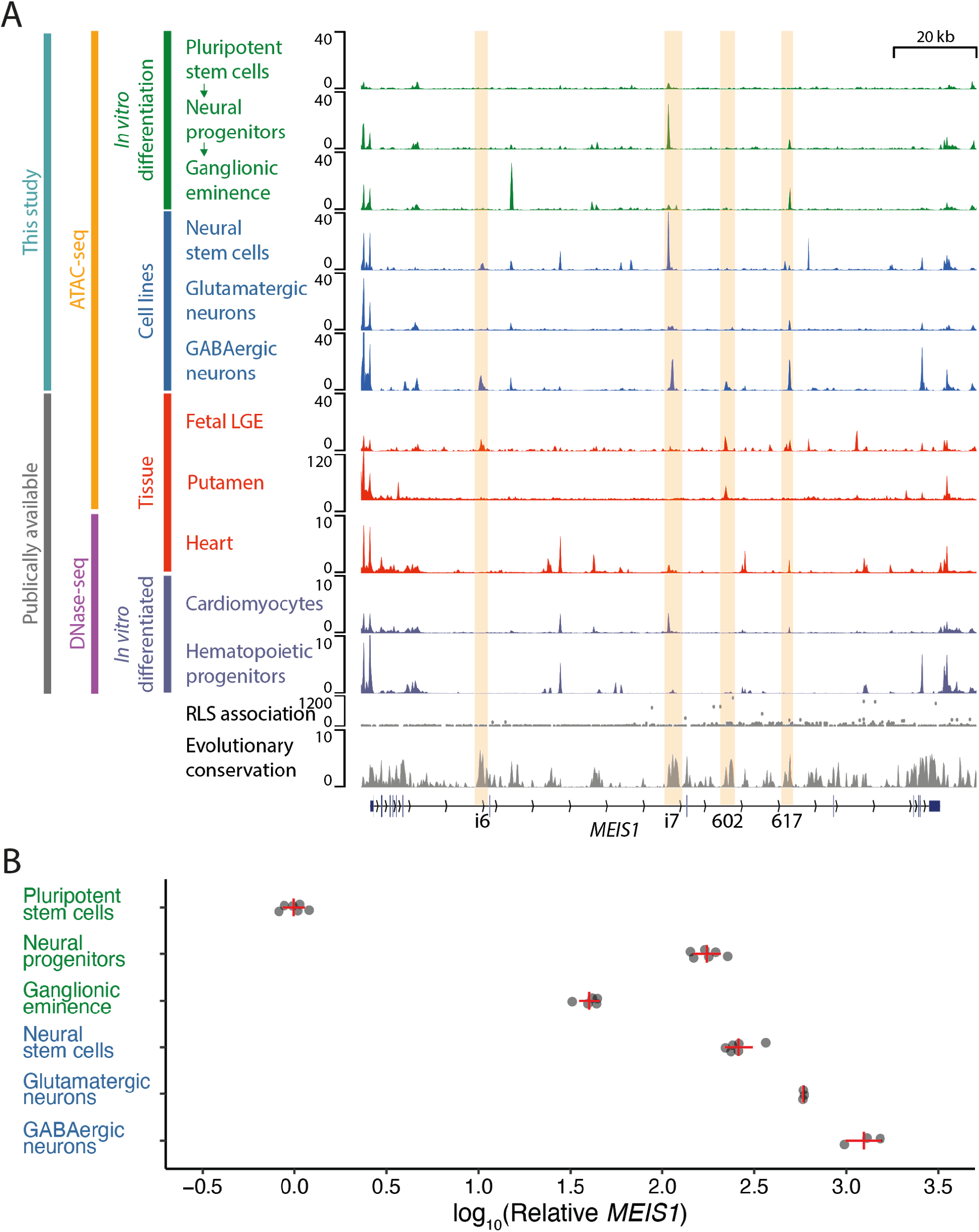
Developmental dynamics of chromatin accessibility in the human *MEIS1* locus. **A**, ATAC-seq (-log_10_P values) and DNase-seq (read depth-normalized signal) of *in vitro* differentiated cells, commercially available cell lines, and tissues. Fetal LGE is 19th gestational week data from (Markenscoff-Papadimitriou et al. 2020). Putamen data are from BOCA (Fullard et al. 2018). Heart, cardiomyocyte, and hematopoietic progenitor data are from ENCODE (ENCODE Project Consortium 2012; Davis et al. 2018). RLS association is from unpublished metaGWAS, scale is -log_10_P. Evolutionary conservation is phyloP 100-way (Pollard et al. 2010). **B**, Relative *MEIS1* expression in the first 6 cell lines depicted in **A**, expressed relative to pluripotent stem cells (=1). Plotted are mean ± SD. The mean expression values ± standard error are: neural progenitors 176 ± 31; ganglionic eminence 40 ± 5; neural stem cells 263 ± 52; glutamatergic neurons 584 ± 6; GABAergic neurons 1,265 ± 277. n=3-6 replicates per cell line.

### Developmental dynamics of chromatin accessibility in the murine *Meis1* locus

In the developing mouse, *Meis1* expression in diverse tissues (limbs, eye, blood, and brain) peaks around mid-gestation (Mercader et al. 1999; Heine et al. 2008; Pineault et al. 2002; Toresson et al. 2000). This is also true of chromatin accessibility (Fig. S2). Moreover, each tissue shows a unique pattern of chromatin accessibility within the *Meis1* locus, with the 617 and 602 elements in particular showing selective accessibility in forebrain, and the i6 element showing selective brain activity (Fig. 4A, Fig. S2). Of note, ChIP-seq analysis at approximately the same embryonic stage revealed that both the 617 and 602 elements are bound by DLX2, a key transcription factor for forebrain GABA neuron differentiation (Panganiban and Rubenstein 2002) (Fig. 4A). Analysis of the temporal dynamics of chromatin accessibility in developing mouse forebrain single nucleus ATAC-seq data support the selective role of element 617 in the inhibitory neuron trajectory (Fig. 4B-D).

**Figure 4.**
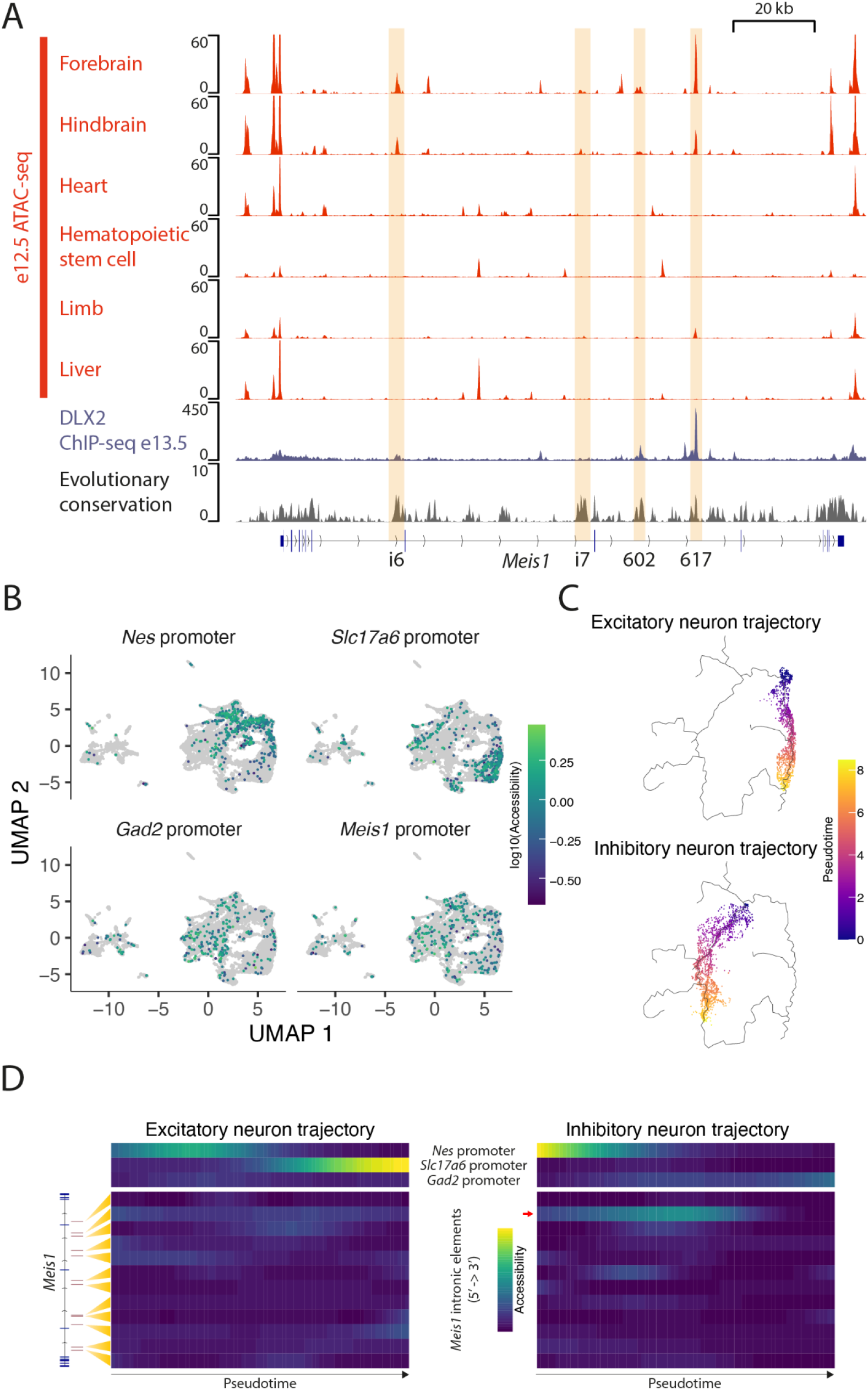
Chromatin accessibility in the murine *Meis1* locus. **A**, ATAC-seq -log_10_P values for 2 replicates of different tissues of embryonic day 12.5 (e12.5) mouse embryos (data from (ENCODE Project Consortium 2012; Davis et al. 2018)); ganglionic eminence DLX2 ChIP-seq at e13.5 (data from (Lindtner et al. 2019)); and evolutionary conservation (phyloP 60-way) in the murine *Meis1* locus, shown here inverted (5’→3’) to compare to the human locus. **B**, Accessibility of promoters for *Nes* (denotes progenitor cells), *Slc17a6* (denotes excitatory neurons), *Gad2* (denotes inhibitory neurons), and *Meis1* in 15,767 single nuclei from developing mouse brain (data from (Preissl et al. 2018)). **C**, Pseudotomporal ordering on uniform manifold approximation and projection (UMAP) coordinates of developing mouse forebrain single nucleus ATAC-seq data in the excitatory (1,484 nuclei) and inhibitory neuron (1,834 nuclei) trajectories. **D**, Pseudotemporal accessibility profiles of three reference promoters and 12 *Meis1* intronic sites in the excitatory and inhibitory neuron trajectories. Element 617 in the inhibitory neuron trajectory is highlighted with a red arrow.

### *MEIS1* enhancer-promoter contacts

Noncoding regulatory elements regulate transcription by establishing direct contacts with gene promoters *via* chromosomal conformational changes (Schoenfelder and Fraser 2019). Intronic elements do not necessarily regulate their host gene, since they can interact topologically with other promoters (Smemo et al. 2014). To determine the topological interactions of the human *MEIS1* promoter, we performed circular chromosome conformation capture followed by sequencing (4C-seq) (Zhao et al. 2006; van de Werken et al. 2012b) in five cell types: induced pluripotent stem cells, neural progenitor cells, ganglionic eminence cells, glutamatergic neurons, and GABAergic neurons (Fig. 5). In all cell types, a gene-spanning contact was detected (Fig. 5), presumably denoting a stable topologically associating domain (TAD) (Nora et al. 2012). In contrast, interaction of element 617 and an additional intron 8 element with the *MEIS1* promoter only reached statistical significance in cell types of the inhibitory neuron trajectory (Fig. 5). The interactions corresponded with the accessibility of intronic elements, in accordance with the functional relatedness of chromatin accessibility and intrachromosomal contacts (Klemm et al. 2019). Supporting our findings, a recently published analysis of cell type-specific chromosome conformation in the developing human cortex (Song et al. 2020) reveals an array of interactions between the *MEIS1* promoter and intronic elements, particularly in radial glia and inhibitory neurons (Fig. S4).

**Figure 5.**
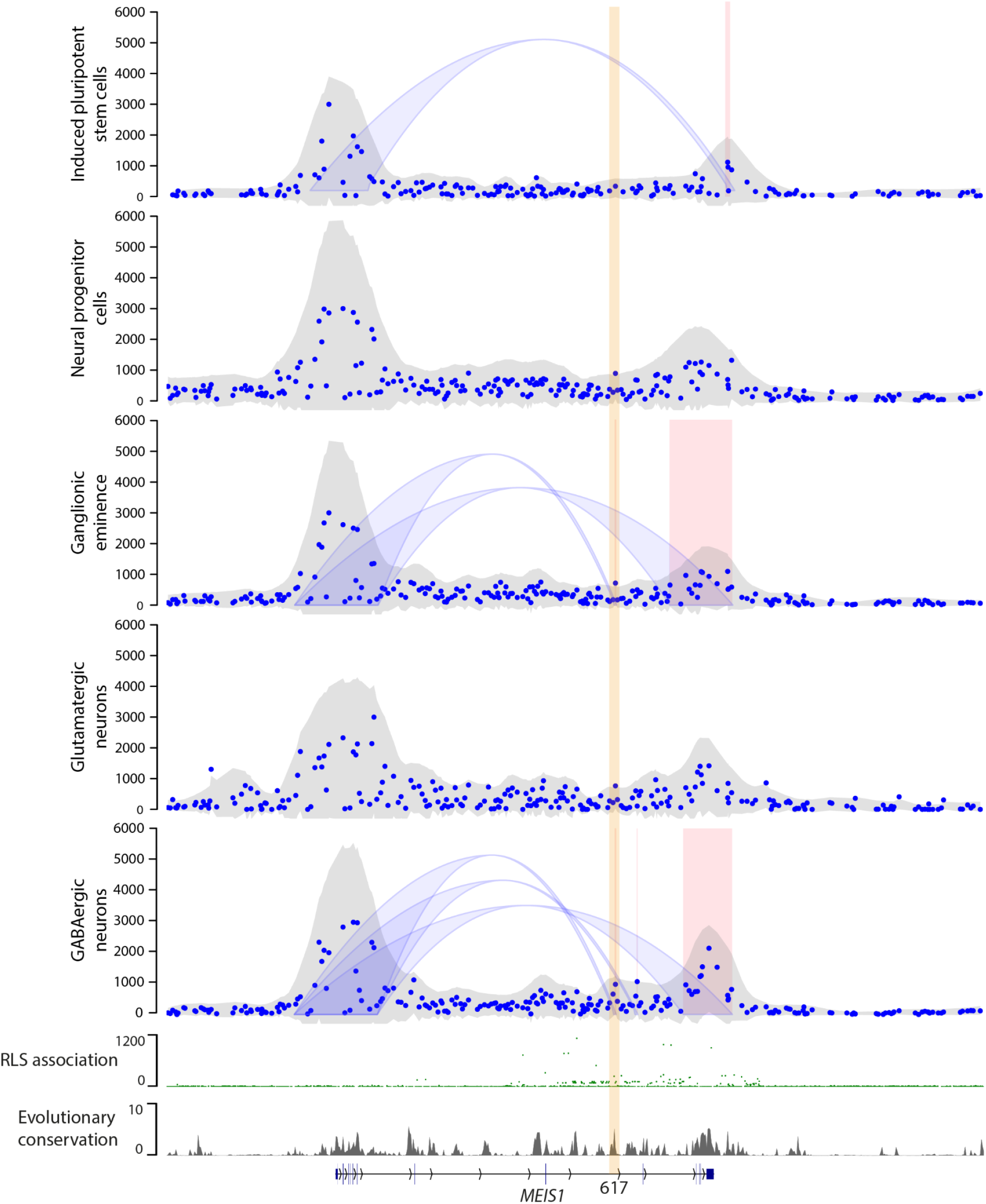
Topological conformation of the *MEIS1* locus. **A**, 4C-seq counts and significant interactions (blue arcs & red rectangles) in human pluripotent cells, neural progenitor cells, ganglionic eminence-like cells, glutamatergic neurons, and GABAergic neurons. RLS association is from unpublished metaGWAS, scale is -log_10_P. Evolutionary conservation is phyloP 100-way.

### Regulation of *MEIS1* expression by enhancer demethylation

To determine the relationship between enhancer methylation and *MEIS1* expression, we characterized methylation using a microarray approach in three cell types: induced pluripotent stem cells (iPSC), neural progenitor cells differentiated from iPSCs with fibroblast growth factor (NPC-FGF), and commercially available neural stem cells (NSC). Neural cells expressed *MEIS1*, while iPSCs had negligible expression (relative to NSCs, mean ± SEM: iPSCs, 0.5 ± 0.02; NSCs 100 ± 3; NPC-FGF 1,467 ± 27; Fig. 6A). Across the locus, *MEIS1*-expressing neural cells had slightly higher levels of methylation. In contrast, elements 617 and i6 were strongly hypomethylated in neural cells (Fig. 6B; cg06919693 within 617, NSC vs iPSC *P*_adj_=6.2×10^−13^, NPC-FGF vs iPSC *P*_adj_=1.3×10^−12^; 6 CpGs within i6, NSC vs iPSC *P*=2.1×10^−24^, NPC-FGF vs iPSC *P*=2.9×10^−47^). We also compared our results to published whole genome bisulfite sequencing results at multiple stages of isogenic *in vitro* neural differentiation, with broadly similar results (Fig. S5). Specifically, the previously published studies (Ziller et al. 2015; Stadler et al. 2011) found moderate demethylation of *MEIS1* intronic elements i6, i7, and 617, particularly at later stages of differentiation (Fig. S5).

**Figure 6.**
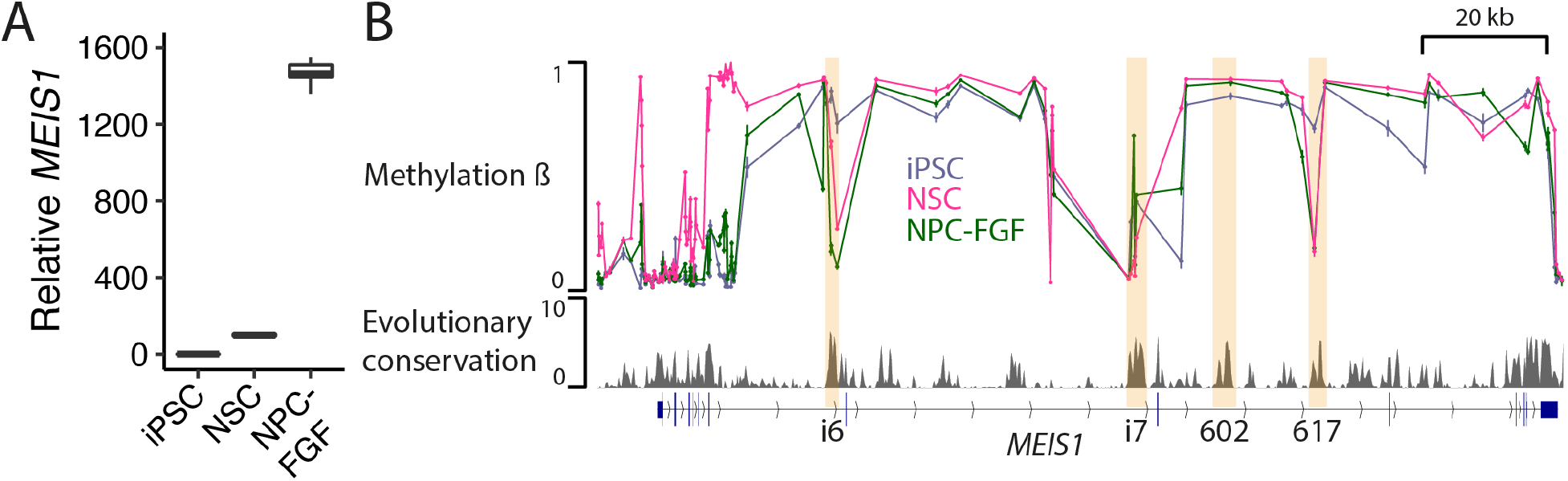
DNA methylation in the *MEIS1* locus. **A**, *MEIS1* expression by qPCR in induced pluripotent stem cells (iPSC), neural stem cells (NSC), and iPSC-derived neural progenitor cells (NPC-FGF). Expression is relative to NSCs (=100) shown as boxplots (n=6 per cell type). **B**, Methylation β values for 123 CpG sites within the *MEIS1* locus for the three cell types (n=6 per cell type). Plotted are mean ± SEM. Also shown is evolutionary conservation (phyloP 100-way).

### *MEIS1* expression quantitative trait loci (eQTLs)

Harnessing large-scale genotype- and tissue-specific gene expression data from GTEx (Consortium 2020), we inspected eQTLs of *MEIS1* expression. eQTLs for different tissues and cell types occurred at vast linear chromosomal distances from the *MEIS1* gene body (extending to ∼550 kb 5’ and ∼830 kb 3’). The eQTLs showed a distinctive tissue-specific distribution, with for example amygdala eQTLs clustering distally 5’ to the gene, cerebellum, tibial nerve, and fibroblast eQTLs distally 3’ to the gene, and lung eQTLs proximally 5’ to and within the gene (Fig. 7A). Of note, caudate and cortex eQTLs clustered in intron 8 and 9, including within elements 602 and 617 (Fig. 7B). As the ganglionic eminences give rise to both caudate (part of the striatum) and cortex, these eQTLs likely relate to effects of intron 8/9 variants that persist through development and into adulthood in cells of the ganglionic eminence lineage. The effect sizes also reflect the proportion of ganglionic eminence-derived GABAergic neurons. Effect sizes in caudate (80-90% of neurons GE-derived GABAergic (Graybiel 1990; Olsson et al. 1998)) were consistently larger than cortex (∼25% of neurons GE-derived GABAergic (Wonders and Anderson 2006)). Overall, *MEIS1* eQTLs show a remarkable tissue-specific segregation within and beyond the locus.

**Figure 7.**
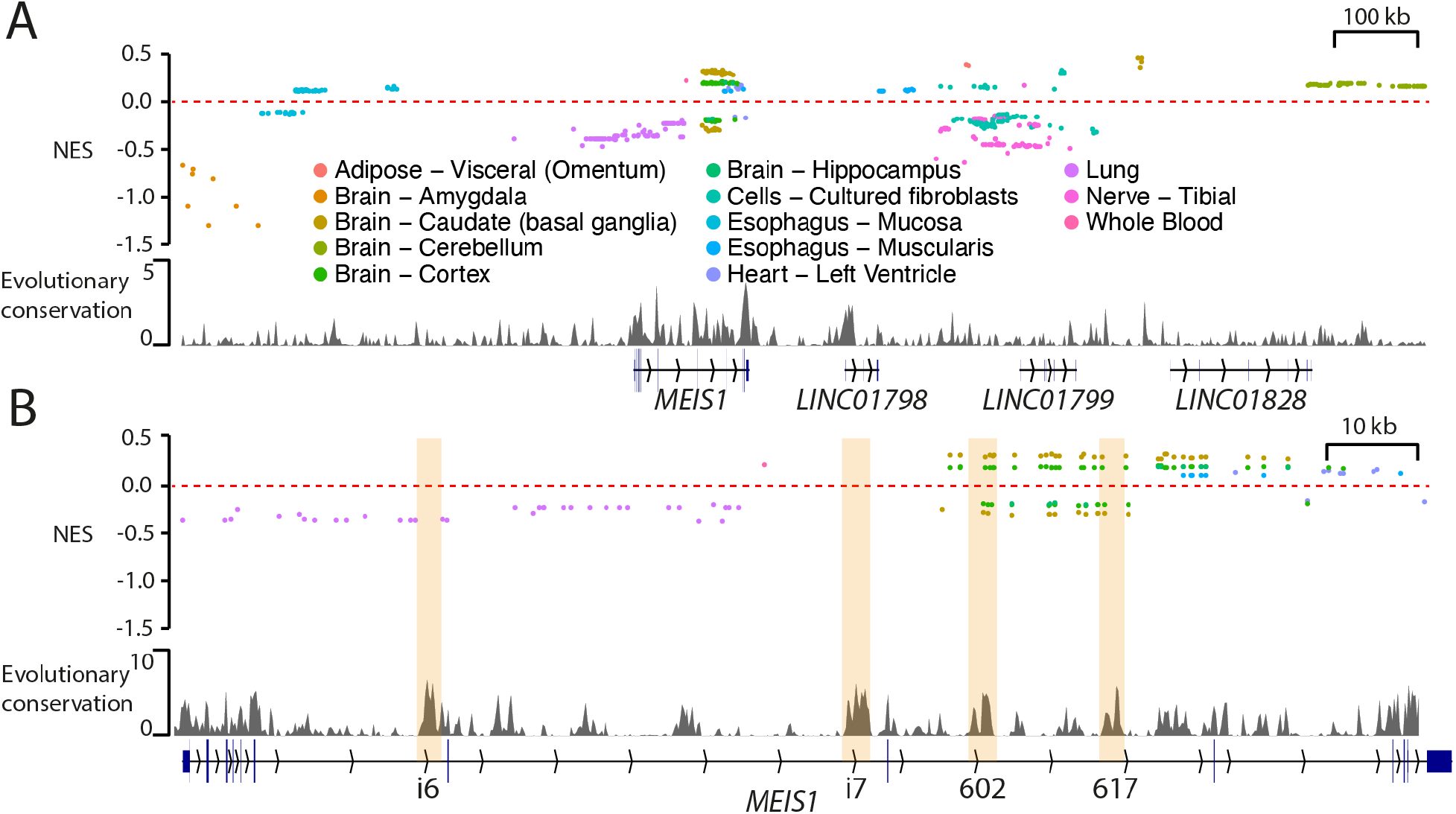
MEIS1 eQTLs. **A**, All 533 *MEIS1* eQTLs from 13 tissues plotted by genomic position and normalized effect size (NES). Evolutionary conservation (phyloP 100-way) is also shown. **B**, 157 *MEIS1* eQTLs mapping within the *MEIS1* locus. The data are identical to **A** at higher genomic resolution.

**Figure 7.**
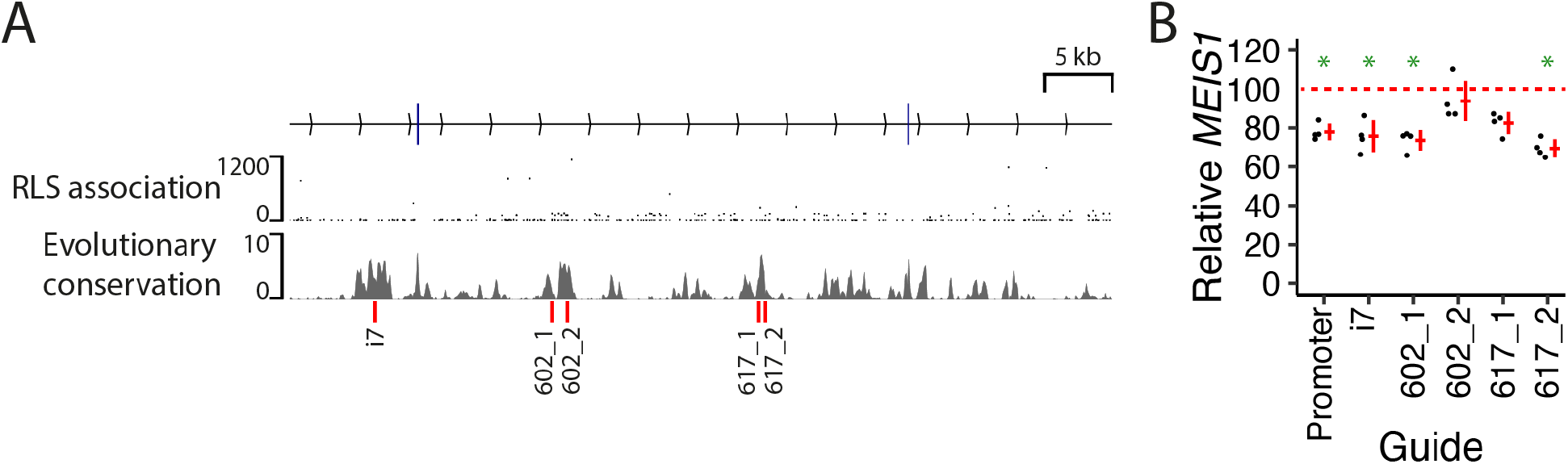
CRISPR interference of intronic *MEIS1* elements. **A**, Genomic location of CRISPR guide target sites (red lines, bottom). Also shown are RLS association (unpublished metaGWAS, scale is -log_10_P) and evolutionary conservation (phyloP 100-way). **B**, *MEIS1* qPCR from NSCs subjected to CRISPR interference with the indicated guides, relative to a *UBQLN2* promoter guide (n=4 replicates per guide). * p < 0.05, one-sided *t*-test compared to *UBQLN2* promoter guide.

### CRISPR interference of intronic *MEIS1* elements reveals regulatory interplay

Several intronic *MEIS1* elements show epigenetic features characteristic of regulatory elements, including chromatin accessibility, contacts with the *MEIS1* promoter, differential methylation, and eQTL effects. However, direct evidence of enhancer activity depends on observing transcriptional effects of functional perturbation. We used CRISPR interference (Gilbert et al. 2013; Thakore et al. 2015; Fulco et al. 2016) to interrogate the functional contribution of three elements to *MEIS1* expression in neural stem cells. We observed an effect of suppression of the intron 7, 602, and 617 elements on *MEIS1* transcription, with a magnitude similar to targeting the *MEIS1* promoter (20-30% suppression of transcription). Thus, in human neural stem cells, these three elements are all involved in regulating *MEIS1* expression, supporting *MEIS1* transcription to a similar extent.

## DISCUSSION

The strong association and comparatively strong effect size of intronic *MEIS1* variants with sleep-related phenotypes (for example, *P* = 1.1 × 10^−180^ and OR = 2 for rs113851554 in RLS metaGWAS (Schormair et al. 2017) indicate a prominent role for this noncoding region in the development of these traits. In this study, we functionally dissected this region to better understand its regulatory function.

We found that specific intronic elements have specific activity, inferred from accessibility, topology, and methylation, with respect to cell types, maturation stage, and differentiation lineage. In particular, element 617 showed a transient peak in activity corresponding to late neurogenesis, and 602 showed persistent activity, particularly in the GABAergic lineage. Element 602 appeared highly selective to this lineage, while element 617 also showed some activity in the cardiogenic and limb development context. In agreement with a particularly important role for elements 617 and 602 in GABAergic differentiation, both are bound by DLX2, a key transcription factor for forebrain GABA neuron differentiation (Panganiban and Rubenstein 2002). Elements i7 and i6 were also active in the GABA neuron lineage to differing degrees of specificity.

There was high concordance of *MEIS1* intronic element accessibility and enhancer-promoter contact frequency in proliferating neural cells, in agreement with the prevailing view that formation of enhancer-promoter contacts and increased enhancer accessibility are mutually reinforcing and functionally related (Klemm et al. 2019; Schoenfelder and Fraser 2019). The enhancer activity corresponding to a maturation stage when cells are transitioning from mitotic to post-mitotic corresponds with the known role of MEIS1 in regulating the cell cycle and balance between proliferation and differentiation (Schulte and Geerts 2019). The interactions between the *MEIS1* promoter and intronic elements in neural development, particularly in the inhibitory neuron lineage, reinforce the importance of *MEIS1* and its regulatory elements in this context.

We also found that neural cells expressing *MEIS1* tended to have higher DNA methylation across the *MEIS1* locus, except at elements 617 and i6. As DNA methylation, and enhancer methylation specifically, typically exert repressive effects on transcription *via* effects on transcription factor binding (Robertson 2005; Aran and Hellman 2013; Yin et al. 2017), we infer that hypomethylation of these elements is permissive for *MEIS1* transcription. Pluripotent cells differentiating into neural cells undergo broad changes in DNA methylation, with demethylation predominating over hypermethylation as genes and regulatory elements supporting differentiation become activated (Ziller et al. 2015; Stadler et al. 2011). Indeed, regions becoming demethylated during neural differentiation are enriched for enhancers near induced genes and their methylation levels are inversely correlated with expression of the nearest gene (Ziller et al. 2015; Stadler et al. 2011). Thus, enhancer demethylation may be an important mechanism of *MEIS1* regulation by facilitating transcription factor binding.

As indicated by eQTL results, the regulation of *MEIS1* in different tissues and cell types is organized in a modular fashion, with regulatory elements for specific tissues clustered in distinct genomic compartments. eQTLs for forebrain tissues cluster clearly within intron 8/9, where the association with sleep-related phenotypes is strongest.

Finally, we show that epigenetic suppression of elements i7, 602, and 617 by targeted CRISPR interference attenuates *MEIS1* expression to a similar extent as suppression of the *MEIS1* promoter in human neural cells. Thus, we assume that each of these elements contributes to a similar extent to the promoter in supporting *MEIS1* transcription. Future combinatorial suppression experiments may provide insight into the extent of synergism between these elements, while suppression experiments in different cell types would enhance understanding of their cell type-specific roles.

In summary, the introns of *MEIS1* are scattered with apparent regulatory elements. Their organization into distinct modules appears to underlie the rich pleiotropy of the gene, with related but distinct functions in diverse tissues and developmental stages. One such module, mostly within intron 8 but extending to neighbouring introns, is prominently associated with sleep/wake-related phenotypes including RLS and insomnia. Here, we show that distinct elements within this module are selectively engaged during differentiation of inhibitory neurons, exerting temporally specific enhancer effects. Genetic variation in *MEIS1* intronic elements probably tunes inhibitory neuron differentiation, thus underlying the association with sleep/wake-related phenotypes.

## METHODS

### Cell lines

The HMGU1 and HMGU12 iPSC lines were obtained from the Helmholtz Center Munich iPSC Core Facility. The lines were generated from the BJ (ATCC CRL-2522) line of foreskin fibroblasts from a healthy newborn human male donor by transfection of six mRNA reprogramming factors (*OCT4, SOX2, KLF4, LIN28, MYC, NANOG;* StemRNA-NM Reprogramming Kit, Stemgent). Cells were maintained in mTeSR culture media (STEMCELL Technologies) on cell culture dishes coated with Matrigel (Corning) or Geltrex (Thermo Fisher).

Human neural stem cells derived from H9 (WA09) human embryonic stem cells were purchased from Thermo Fisher. The cells were cultured in complete StemPro NSC SFM (Thermo Fisher) consisting of KnockOut D-MEM/F-12 with StemPro Neural Supplement, EGF, bFGF, and GlutaMAX on Geltrex-coated cell culture dishes.

GABAergic neurons and glutamatergic neurons (iCell GABANeurons Kit and iCell GlutaNeurons Kit) were purchased from Cellular Dynamics. iCell GABANeurons and GlutaNeurons are highly pure populations of human neurons, comprised primarily of GABAergic and cortical glutamatergic neurons respectively, derived from induced pluripotent stem (iPS) cells using proprietary differentiation and purification methods. Cells were collected for downstream experiments five days post-plating.

### *In vitro* differentiation

iPSCs were differentiated towards ganglionic eminence using the protocol from Close et al. (Close et al. 2017) with slight modifications. iPSCs were grown to 90% confluence and then dissociated with TrypLE Select Enzyme for 5 minutes at 37°C. 500,000 iPS cells were plated per well of a 24 well plate in mTeSR supplemented with RevitaCell (Thermo Fisher). The next day (day 1), the medium was replaced with neural induction medium (NIM) consisting of DMEM/F12, N-2, B-27, and GlutaMAX supplements, MEM Non-Essential Amino Acids, 0.11 mM 2-mercaptoethanol, 0.05% (v/v) Bovine Serum Albumin Fraction V Solution, Penicillin-Streptomycin (all Thermo Fisher), 100 nM LDN193189 and 10 μM SB431542 (Cayman Chemical), and 2 μM XAV939 (R & D Systems). NIM was changed daily until day 5, when it was replaced with 75% NIM/25% N2 medium (DMEM/F12, N-2 supplement, 0.15% (w/v) dextrose, 55 μM 2-mercaptoethanol, Penicillin-Streptomycin). On day 7, cells were fed with 50% NIM/50% N2, and on day 9 with 25% NIM/75% N2. On day 10, cells were dissociated into a single cell suspension with TrypLE Express Enzyme and plated onto Matrigel-coated 24-well plates in 25% NIM/75% N2 supplemented with RevitaCell at 2 million cells per well. At this stage some material was harvested for RNA extraction and ATAC-seq (NPC stage). On day 11 the medium was replaced with N2/B27 medium (N2 medium with B-27 supplement) containing 0.65 µM purmorphamine (Cayman Chemical). On day 19-23, cells were fed daily with N2/B27 medium without purmorphamine. On day 24, cells were collected for RNA extraction and ATAC-seq.

To generate NPC-FGF cells used for DNA methylation studies, the protocol from Reinhardt et al. was used (Reinhardt et al. 2013). iPSCs were dissociated with 2 mg/ml collagenase IV (Thermo Fisher) for 45 min at 37°C, which was then neutralized with mTeSR. The cells were centrifuged at 200 g for 4 min, then resuspended in differentiation medium (20% KnockOut Serum Replacement and MEM Non-Essential Amino Acids in DMEM-F12, all Thermo Fisher) supplemented with a small molecule cocktail (1 µM dorsomorphin, 10 µM SB431542, 3 µM CHIR99021, and 0.5 µM purmorphamine) and RevitaCell, and grown in ultra low attachment plates to form embryoid bodies. The medium was replaced the following day (day 2). On day 3, the medium was changed to N2B27 medium (DMEM-F12 and Neurobasal 1:1, B-27 and N-2 supplements, and GlutaMAX) with the small molecule cocktail. The medium was changed on day 4, and on day 5 without purmorphamine and CHIR99021. On day 6, embryoid bodies were plated on Geltrex-coated plates in N2B27 medium with dorsomorphin, SB431542, and basic fibroblast growth factor (bFGF, 10 ng/ml). This medium was replaced daily. Cells were propagated for 3 passages after Matrigel plating, then harvested at confluence.

### ATAC-seq

We used the previously described protocol of Buenruostro *et al*. (Buenrostro et al. 2015) with 50,000 cells per replicate. Adherent cells were dissociated with TrypLE Select Enzyme and centrifuged at 500 g for 5 min at 4°C. The cell pellet was resuspended in ice-cold lysis buffer (10 mM Tris-HCl, pH 7.4, 10 mM NaCl, 3 mM MgCl_2_, 0.1% IGEPAL CA-630) and centrifuged for 20 min at 500 g at 4°C. The pellet was resuspended in transposition reaction mix (Nextera Tn5 transposase, Illumina) and incubated for 30 min at 37°C. Samples were purified using the MinElute Kit (Qiagen).

Purified samples were then amplified by PCR (maximum 12 cycles) using barcoded primers from the Nextera Index Kit (Illumina). To determine the optimal number of cycles for each sample, samples were first amplified for 5 cycles (5 min at 72°C, 30 sec at 98°C, and 5 cycles of 10 sec at 98°C, 30 sec at 63°C, and 1 min at 72°C). Samples amplified for 5 cycles were then subjected to qPCR (SYBR Green, Thermo Fisher). The additional number of cycles required was the number of cycles required to reach one quarter of the maximal fluorescence intensity in the first 5 cycles. The remainder of each sample was subsequently amplified for an additional 5-7 cycles.

Amplified samples were purified with SPRI beads (Beckman Coulter) at a ratio 1:1.8 sample:beads, and fragment size was checked using the High Sensitivity DNA kit (Agilent). Libraries were sequenced on a HiSeq 4000 (Illumina), two samples per lane, 100 bp paired end reads. Reads were trimmed and aligned with bowtie2 (Langmead and Salzberg 2012), with standard parameters and a maximum fragment length of 2,000. Duplicate reads were removed with Picard. De-duplicated reads were filtered for high quality (samtools (Li et al. 2009), MAPQ ≥ 30), nonmitochondrial chromosome, non-Y chromosome, and proper pairing (samtools flag 0 × 2). Peaks were called with macs2 (Zhang et al. 2008, 2)], and filtered out with IDR threshold of 0.1 (Li et al. 2011) and blacklist of artifactual regions in hg19. Libraries were quality controlled by downsampling to 5 million reads and evaluating transcription start site enrichment (RefSeq) and fraction of reads in peaks.

### Single nucleus ATAC-seq analysis

Single nucleus ATAC-seq data from developing mouse forebrain (Preissl et al. 2018) were downloaded from GEO (accession GSE100033). Data were processed with Cicero (Pliner et al. 2018) using default parameters. The trajectory root was chosen manually based on accessibility of the *Nes* promoter.

### 4C-seq

We used the previously described protocol of van de Werken *et al*. (van de Werken et al. 2012a). Cells were dissociated to single cells with TrypLE Express and fixed in 2% formaldehyde in DPBS (Thermo Fisher) for 10 min. Glycine was added to a final concentration of 222 mM to quench the cross-linking reaction. Cells were centrifuged for 10 minutes at 400g at 4°C, the supernatant discarded, and the cell pellet resuspended in cold lysis buffer (50 mM Tris pH 7.5, 150 mM NaCl, 5 mM EDTA, 0.5% NP-40, 1% Triton X-100, cOmplete protease inhibitor cocktail, Roche) and incubated on ice for 30 minutes. To ensure complete lysis, cells were stained with methyl green pyronin (Dianova). In the case of efficient lysis, nuclei are stained blue and cytoplasm pink. Next, lysed cells were centrifuged and the pellet was washed with DPBS and resuspended in water. NEBuffer and SDS (0.3% final concentration) were added and the samples were incubated at 37°C for 1 hr while shaking at 900 rpm with occasional pipetting to break cell aggregates. Triton X-100 (Biomol) was added to a concentration of 2.5% and samples were again incubated at 37°C for 1 hr while shaking.

The first round of digestion was done with DpnII (New England Biolabs) at 37°C overnight. Digest efficiency was checked on an agarose gel. DpnII was inactivated by incubating for 20 min at 65°C. Ligation was performed with T4 ligase (Thermo Fisher) overnight at 16°C and checked on an agarose gel. Samples were then decrosslinked with Proteinase K at 65°C overnight. After DNA purification with phenol-chloroform, the second round of digestion was performed with Csp6I (Thermo Fisher). Digestion was followed by overnight ligation with T4 ligase at 16°C. Samples were purified using Amicon Ultra-15 columns (Millipore) and DNA quantified using the Qubit dsDNA HS Assay Kit (Thermo Fisher).

To generate 4C libraries, we used 3.2 µg of DNA for PCR amplification divided into 16 PCR reactions, using Expand Long Template PCR System (Sigma Aldrich). PCR primers were designed using the *MEIS1* promoter as a viewpoint and included Truseq adapters. Primer genomic targets are shown in Fig. S3 and sequences in Table S1. PCR conditions were 2 min at 94°C, 30 cycles of 15 sec at 94°C, 1 min at 55°C, and 3 min at 68°C followed by 5 min at 68°C. Pooled libraries were purified using SPRI beads (Beckman Coulter) at a ratio of 1:1.8 sample:beads, and fragment size was checked using the High Sensitivity DNA kit (Agilent). The libraries were sequenced on an Illumina MiSeq. Reads were trimmed with Trimmomatic v0.36 (Bolger et al. 2014), aligned with bowtie2 (Langmead and Salzberg 2012) and subsequently processed, filtered, analyzed and visualized according to Basic4Cseq (Walter et al. 2014) in R v 4.0.2.

### DNA methylation analysis

Genome-wide DNA methylation from induced pluripotent stem cells (iPSC), neural stem cells (NSC), and iPSC-derived neural progenitor cells (NPC-FGF) was assessed using the Infinium Human Methylation EPIC BeadChip (Illumina), according to the manufacturer’s protocol. The EPIC array covers over 850,000 methylation sites. Genome-wide raw signal intensity data were subjected to a standard quality control (QC) and quantile normalization pipeline using the minfi (Aryee et al. 2014) and limma (Ritchie et al. 2015) packages in R v 4.0.2. Probes with poor quality (<95% call rate or detection *P* value > 0.01), SNPs at the CpG site, and cross-reactivity were filtered out prior to differential methylation analysis. Methylation level for each CpG site was calculated as a beta value β=M/(M+U+100), where M>0 and U>0 denote the methylated and unmethylated signal intensities measured by the array, and the offset of 100 is added to M+U to stabilize beta values when both M and U are small. Beta values range between 0 and 1, with 0 indicating no methylation and 1 full methylation. CpG sites were classified as hypo- or hypermethylated according to ENCODE guidelines. Probe-wise and region-wise differential methylation between the different cell types were estimated using M values in limma, obtaining moderated t-statistics, *P* values and adjusted *P* values using false discovery rate (FDR) for each CpG site.

### eQTL analysis

*MEIS1* eQTL data were downloaded from GTEx (Consortium 2020) and plotted with ggplot2 (Wickham 2016, 2) and pyGenomeTracks (Ramírez et al. 2018).

### CRISPR interference

To prepare CRISPRi lentiviruses, Lenti-X™ 293T Cells (Takara) were seeded on 10 cm dishes (6 × 10^6^ cells per dish). The next day, cells were transfected with second generation lentiviral packaging plasmids (pMD2.G, 0.72 pmol; and psPAX2, 1.3 pmol) and the CRISPRi transfer plasmid (pLV hU6-sgRNA hUbC-dCas9-KRAB-T2a-Puro, 1.64 pmol) carrying the respective guide RNA (see Table S1). pMD2.G was a gift from Didier Trono (Addgene plasmid # 12259; http://n2t.net/addgene:12259 ; RRID:Addgene_12259). psPAX2 was a gift from Didier Trono (Addgene plasmid # 12260 ; http://n2t.net/addgene:12260 ; RRID:Addgene_12260). pLV hU6-sgRNA hUbC-dCas9-KRAB-T2a-Puro was a gift from Charles Gersbach (Addgene plasmid # 71236 ; http://n2t.net/addgene:71236 ; RRID:Addgene_71236). Plasmids were mixed with 48 µg polyethylenimine ‘Max’, MW 40,000 Da (Polysciences) in 600 µl OptiMEM (ThermoFisher), incubated for 20 min at room temperature, then added to the Lenti-X cells. Cells were incubated at 37°C for 16 hr, then medium was changed. 48 hr later, the medium was harvested, centrifuged at 1,200 x g for 5 min at 4°C, and filtered through a 0.45 µm polyethersulfone (PES) syringe filter. The filtered supernatant was used to transduce NSCs.

NSCs were seeded in Geltrex-coated 24-well plates (50,000 cells per well). The following day, CRISPRi lentiviral supernatants (100 µl per well) were added. Two days later, cells were selected with puromycin (1 µg/ml) for 24 hr, after which the medium was replaced without puromycin. RNA was harvested the following day (see Quantitative PCR below).

### Quantitative PCR

Total RNA was extracted with the RNeasy Mini Kit (Qiagen). RNA was reverse transcribed into complementary DNA (cDNA) using the High-Capacity cDNA Reverse Transcription Kit (Thermo Fisher). Quantitative PCR was performed in duplicate using TaqMan Universal PCR Master Mix (Thermo Fisher) with TaqMan gene expression assays: *MEIS1* (Hs00180020_m1), *PAX6* (Hs00240871_m1), *FOXG1* (Hs01850784_s1), *GSX2* (Hs00370195_m1), *NKX2-1* (Hs00968940_m1), and *GAPDH* (Hs02758991_g). qPCR reactions were performed on a 7900HT Fast Real-Time PCR System (Applied Biosystems) as follows: 2 min at 50°C, 10 min at 95°C, and 40 cycles of 15 sec at 95°C and 1 min at 60°C. For quantification, the 2^-ΔΔCT^ method was used with *GAPDH* as the endogenous reference.

## DATA ACCESS

All raw and processed sequencing data generated in this study have been submitted to the NCBI Gene Expression Omnibus (GEO; https://www.ncbi.nlm.nih.gov/geo/) under accession number GSE174727.

## COMPETING INTEREST STATEMENT

The authors declare that they have no competing interests

## ACKNOWLEDGEMENTS

The work was supported by funding from the German Federal Ministry for Education and Resarch (BMBF), the French Agence Nationale de la Recherche, and the European Commission (ERA-NET NEURON, “SMART”), as well as the Munich Cluster for Systems Neurology (SyNergy). A.A.N. was supported by a Bayhost Scholarship.

We thank Monika Zimmermann, Julia Vandrey, and Irmgard Zaus for technical assistance. We thank the ENCODE consortium for genomic data, particularly the Bing Ren laboratory for mouse embryonic ATAC-seq data and the John Stamatoyannopoulos laboratory for human DNase-seq data. The Genotype-Tissue Expression (GTEx) Project was supported by the Common Fund of the Office of the Director of the National Institutes of Health, and by NCI, NHGRI, NHLBI, NIDA, NIMH, and NINDS. The data used for the analyses described in this manuscript were obtained from the GTEx Portal on 16 October 2020.

## Author’s contributions

D.D.L. and A.A.N. designed the study and analyzed the data. A.A.N. performed the experiments. D.D.L. wrote the manuscript and prepared the figures with input from all authors. C.Z. and N.M-S. performed bioinformatic analysis. W.K. and J.W. supervised the project. All authors read and approved the final manuscript.

## SUPPLEMENTAL INFORMATION

**Figure S1.**
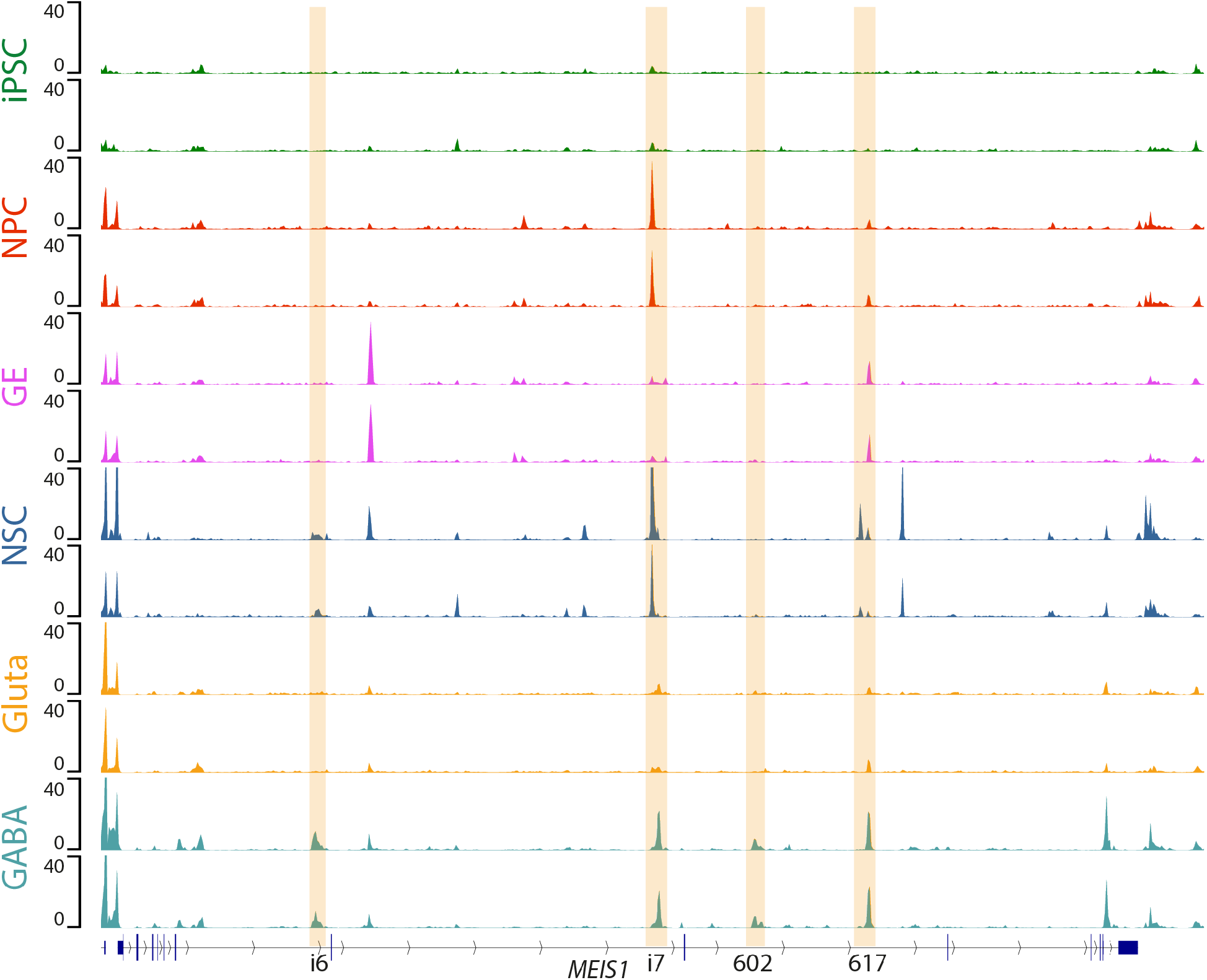
ATAC-seq replicate concordance. ATAC-seq for each cell type was performed in duplicate. Both replicates are shown. Values are -log_10_P.

**Figure S2.**
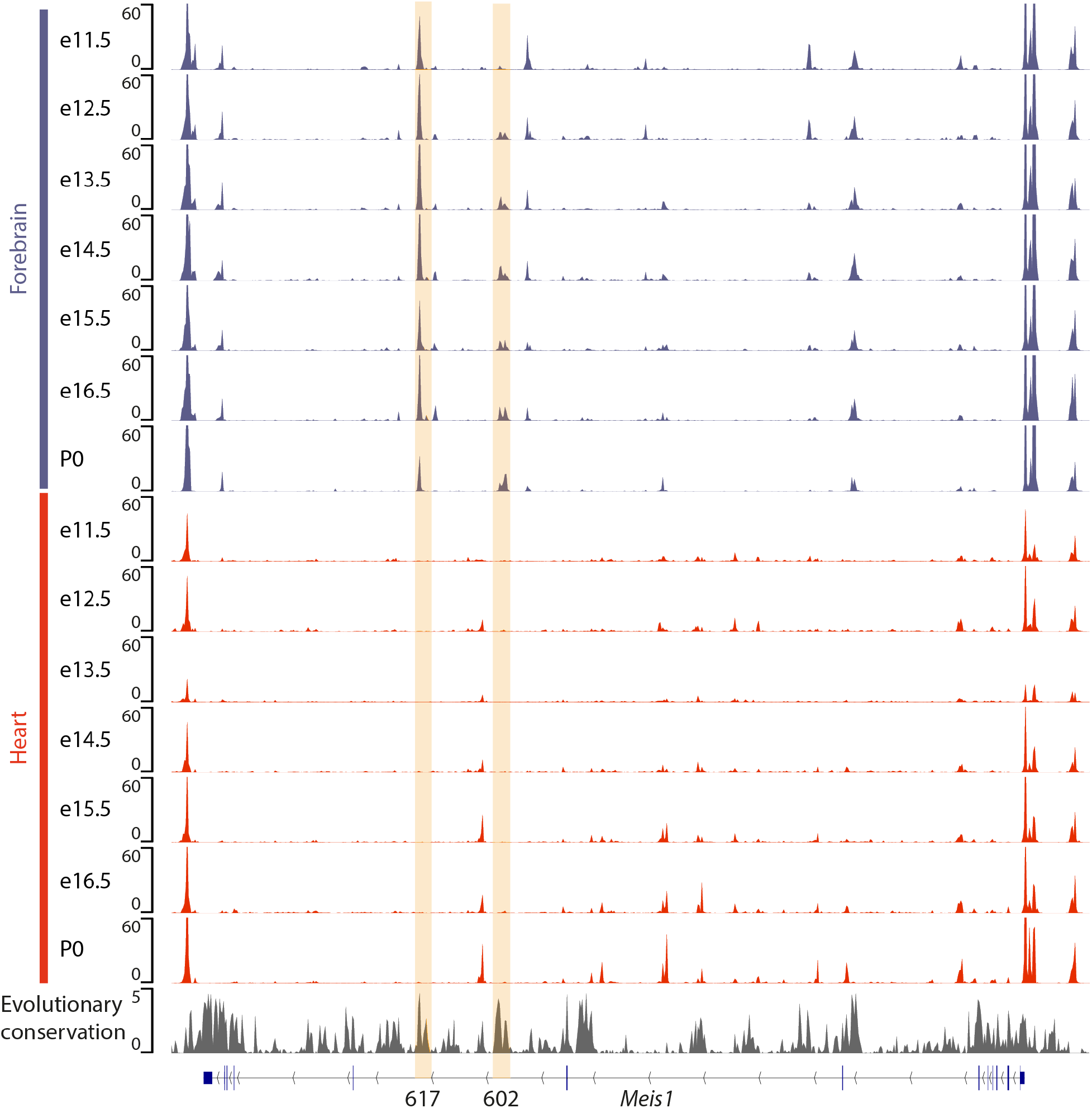
Mouse tissue ATAC-seq at different developmental stages. ATAC-seq signal p-values for 2 replicates of developing mouse forebrain and heart at the indicated developmental stages. Data from ENCODE (ENCODE Project Consortium 2012; Davis et al. 2018). Evolutionary conservation is phyloP 60-way.

**Figure S3.**
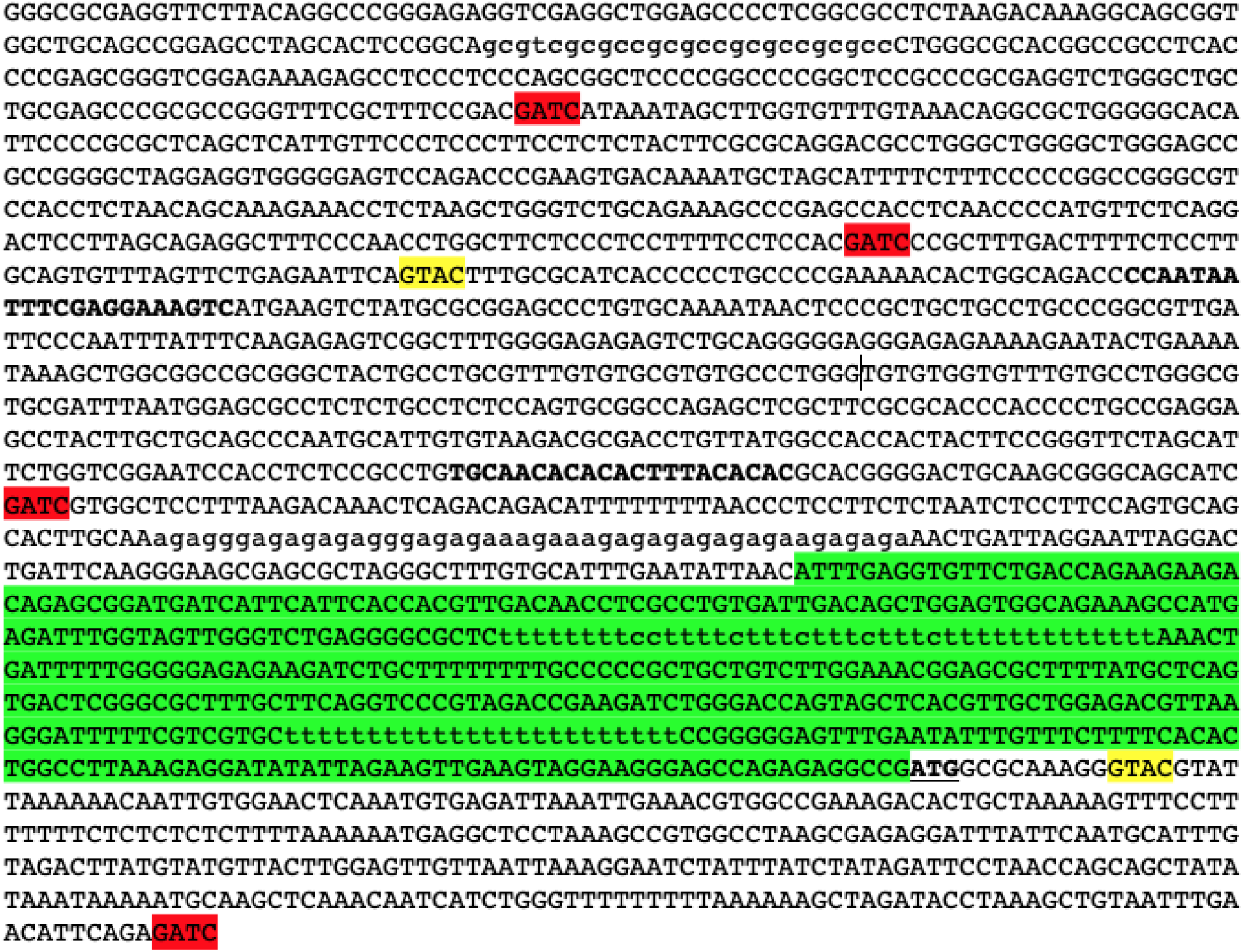
4C-seq primer binding sites. DNA sequence around the human *MEIS1* promoter annotated with 5’ untranslated region (green highlight), *MEIS1* start codon (underlined), DpnII and Csp6I restriction sites (red and yellow respectively), and primer binding sites (bold).

**Figure S4.**
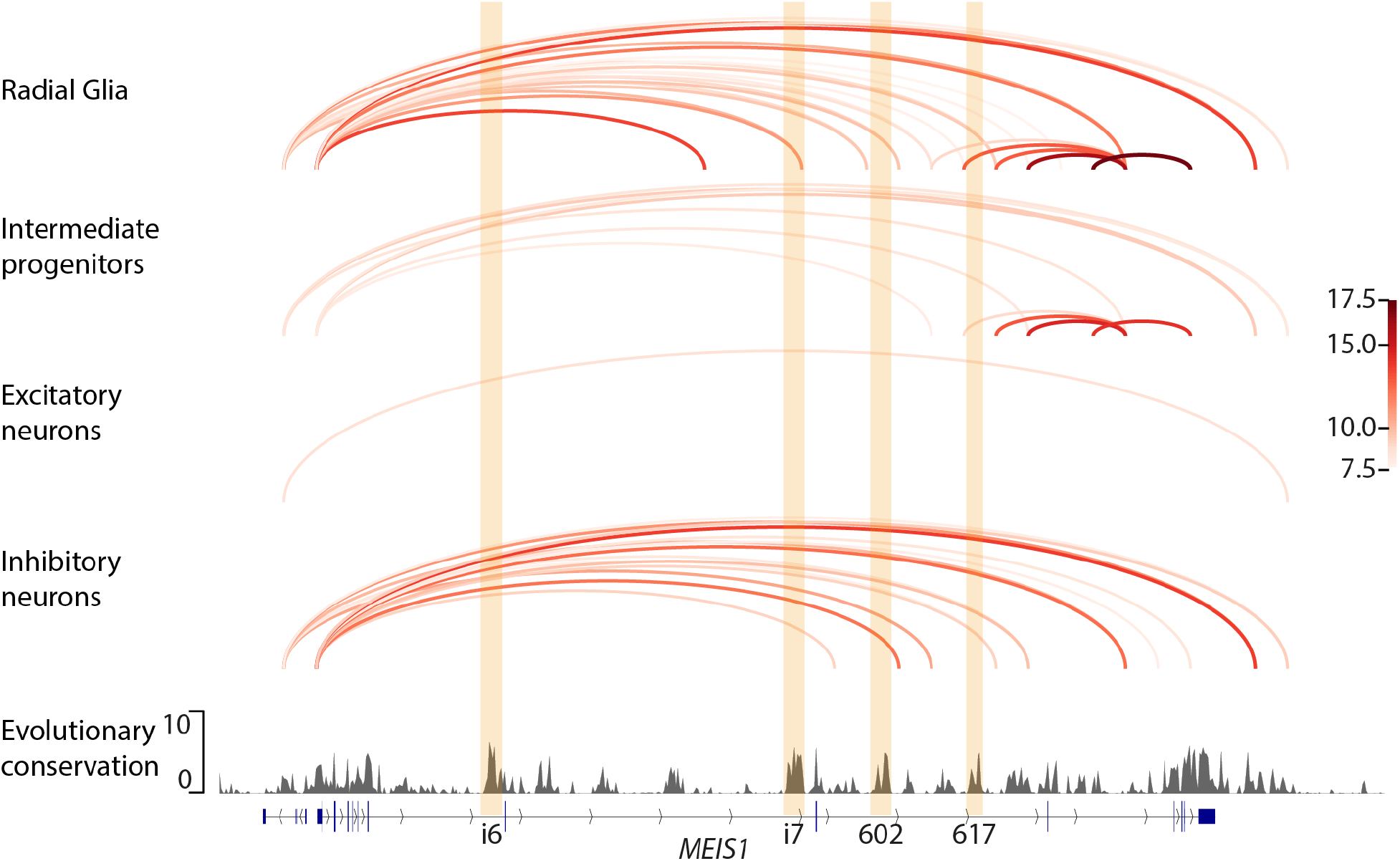
Cell type-specific chromatin interactions in the developing human cortex. Results of proximity ligation-assisted chromatin immunoprecipitation sequencing (PLAC-seq) in four cell types of the developing human cortex from (Song et al. 2020). Arcs represent interactions at a resolution of 5 kb, with colour scale indicating the MAPS -log_10_FDR (Juric et al. 2019).

**Figure S5.**
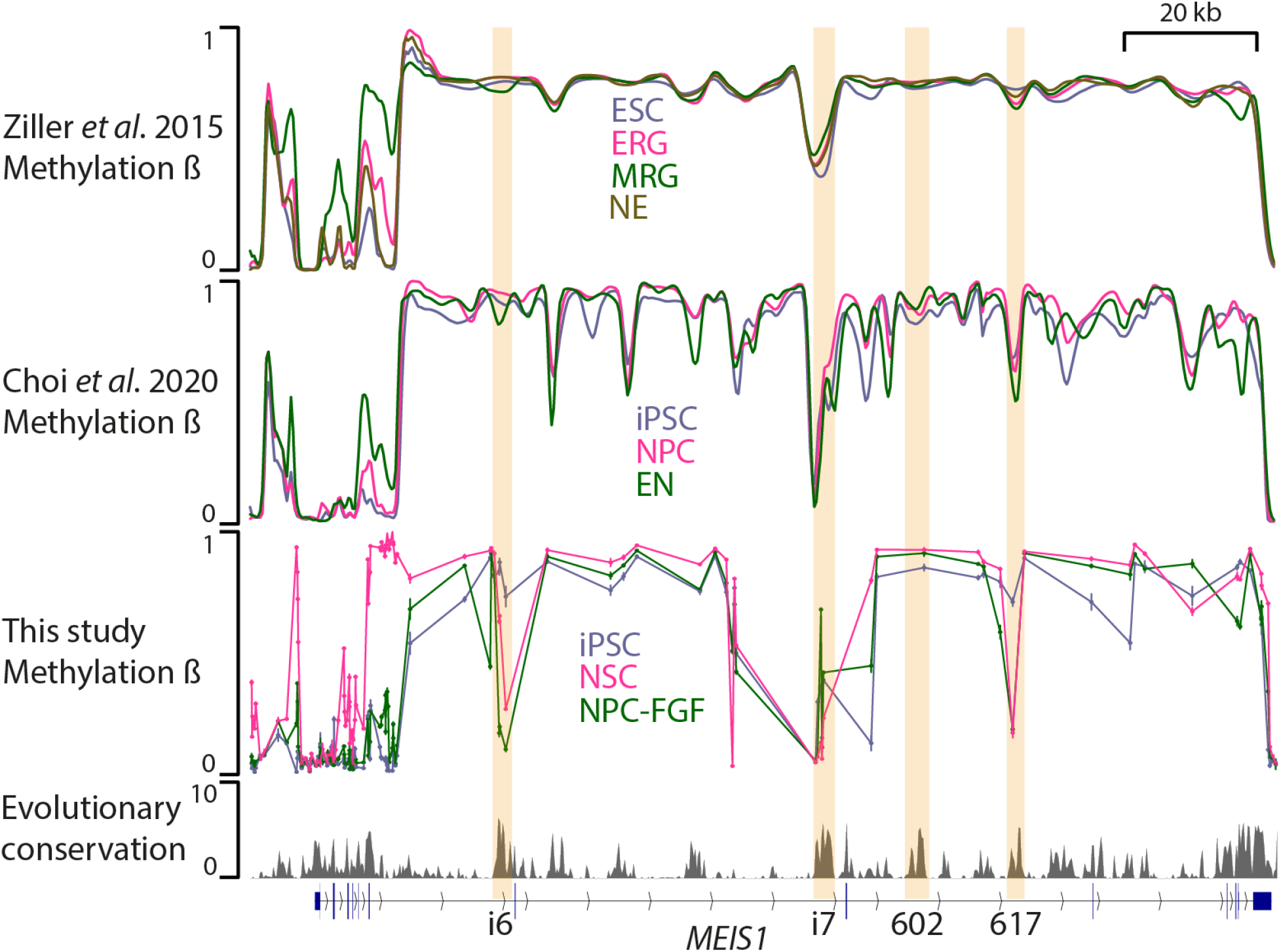
Comparison of *MEIS1* locus methylation profiles with other studies. We compared DNA methylation profiles obtained in our study (third row) with those obtained by whole genome bisulfite sequencing at different stages of isogenic *in vitro* neural differentiation. Data in top row are from (Ziller et al. 2015); data in second row are from (Choi et al. 2020). EN, early neurons; ESC, embryonic stem cells; ERG, early radial glia; iPSC, induced pluripotent stem cells; MRG, mid radial glia; NE, neuroepithelial cells; NPC, neural progenitor cells; NPC-FGF, neural progenitor cells differentiated with fibroblast growth factor; NSC, neural stem cells.

**Table S1.**
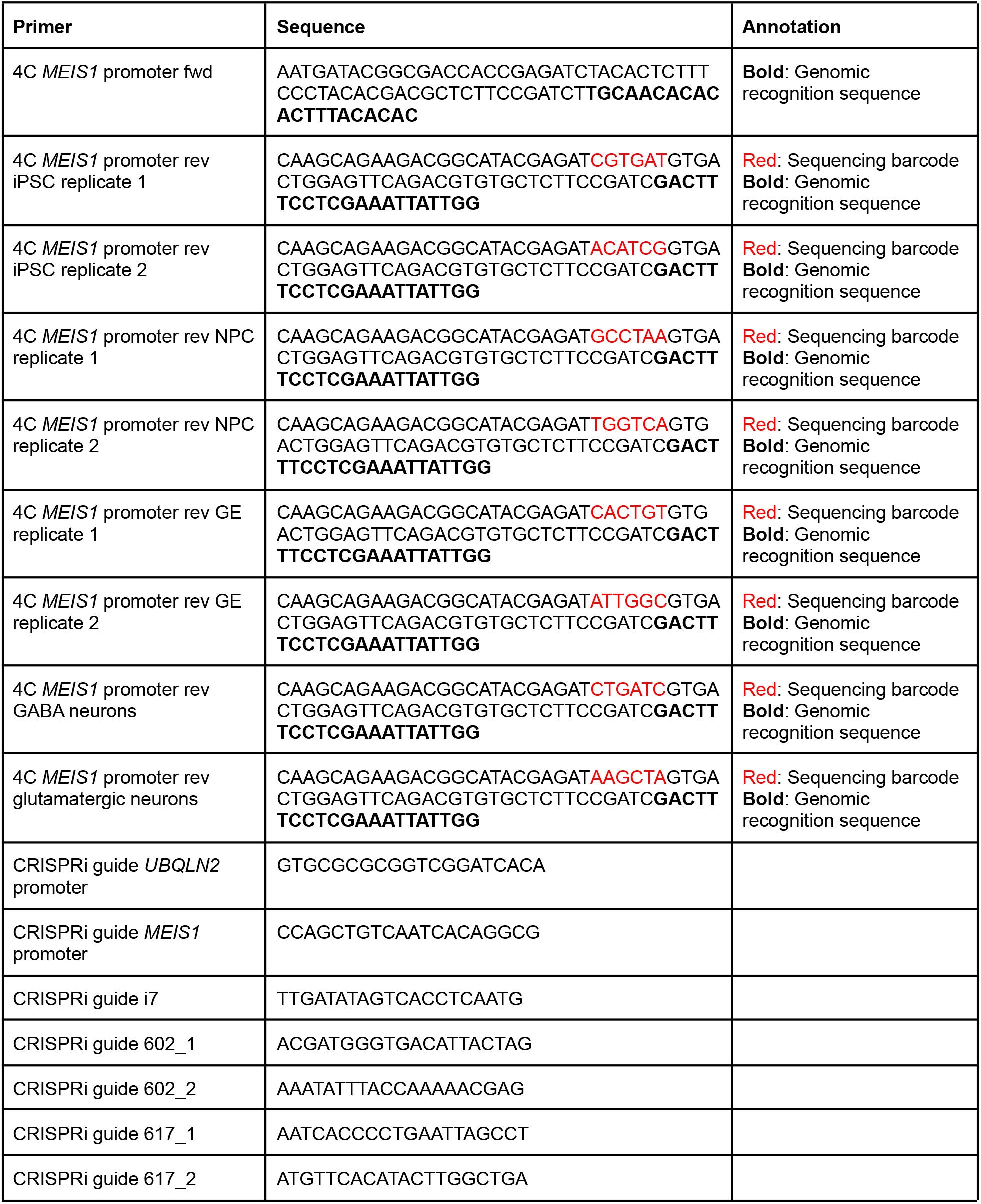
Oligonucleotide sequences.

## Notes

https://www.ncbi.nlm.nih.gov/geo/query/acc.cgi?acc=GSE174727

### Competing Interest Statement

The authors have declared no competing interest.

## REFERENCES

Aran D, Hellman A. 2013. DNA Methylation of Transcriptional Enhancers and Cancer Predisposition. Cell 154: 11–13.

Aryee MJ, Jaffe AE, Corrada-Bravo H, Ladd-Acosta C, Feinberg AP, Hansen KD, Irizarry RA. 2014. Minfi: a flexible and comprehensive Bioconductor package for the analysis of Infinium DNA methylation microarrays. Bioinformatics 30: 1363–1369.

Bolger AM, Lohse M, Usadel B. 2014. Trimmomatic: a flexible trimmer for Illumina sequence data. Bioinformatics 30: 2114–2120.

Buenrostro JD, Wu B, Chang HY, Greenleaf WJ. 2015. ATAC-seq: A Method for Assaying Chromatin Accessibility Genome-Wide. Curr Protoc Mol Biol 109: 21.29.1-21.29.9.

Choi W-Y, Hwang J-H, Cho A-N, Lee AJ, Jung I, Cho S-W, Kim LK, Kim Y-J. 2020. NEUROD1 Intrinsically Initiates Differentiation of Induced Pluripotent Stem Cells into Neural Progenitor Cells. Mol Cells 43: 1011–1022.

Close JL, Yao Z, Levi BP, Miller JA, Bakken TE, Menon V, Ting JT, Wall A, Krostag A-R, Thomsen ER, et al. 2017. Single-Cell Profiling of an In Vitro Model of Human Interneuron Development Reveals Temporal Dynamics of Cell Type Production and Maturation. Neuron 93: 1035–1048.e5.

Consortium TGte. 2020. The GTEx Consortium atlas of genetic regulatory effects across human tissues. Science 369: 1318–1330.

Corbin JG, Gaiano N, Machold RP, Langston A, Fishell G. 2000. The Gsh2 homeodomain gene controls multiple aspects of telencephalic development. Development 127: 5007–5020.

Davis CA, Hitz BC, Sloan CA, Chan ET, Davidson JM, Gabdank I, Hilton JA, Jain K, Baymuradov UK, Narayanan AK, et al. 2018. The Encyclopedia of DNA elements (ENCODE): data portal update. Nucleic Acids Res 46: D794–D801.

Domcke S, Hill AJ, Daza RM, Cao J, O’Day DR, Pliner HA, Aldinger KA, Pokholok D, Zhang F, Milbank JH, et al. 2020. A human cell atlas of fetal chromatin accessibility. Science 370.

ENCODE Project Consortium. 2012. An integrated encyclopedia of DNA elements in the human genome. Nature 489: 57–74.

Fulco CP, Munschauer M, Anyoha R, Munson G, Grossman SR, Perez EM, Kane M, Cleary B, Lander ES, Engreitz JM. 2016. Systematic mapping of functional enhancer–promoter connections with CRISPR interference. Science 354: 769–773.

Fullard JF, Hauberg ME, Bendl J, Egervari G, Cirnaru M-D, Reach SM, Motl J, Ehrlich ME, Hurd YL, Roussos P. 2018. An atlas of chromatin accessibility in the adult human brain. Genome Res 28: 1243–1252.

Gilbert LA, Larson MH, Morsut L, Liu Z, Brar GA, Torres SE, Stern-Ginossar N, Brandman O, Whitehead EH, Doudna JA, et al. 2013. CRISPR-mediated modular RNA-guided regulation of transcription in eukaryotes. Cell 154: 442–451.

Graybiel AM. 1990. Neurotransmitters and neuromodulators in the basal ganglia. Trends Neurosci 13: 244–254.

Hammerschlag AR, Stringer S, de Leeuw CA, Sniekers S, Taskesen E, Watanabe K, Blanken TF, Dekker K, Te Lindert BHW, Wassing R, et al. 2017. Genome-wide association analysis of insomnia complaints identifies risk genes and genetic overlap with psychiatric and metabolic traits. Nat Genet 49: 1584–1592.

Heine P, Dohle E, Bumsted-O’Brien K, Engelkamp D, Schulte D. 2008. Evidence for an evolutionary conserved role of homothorax/Meis1/2 during vertebrate retina development. Development 135: 805–811.

Jansen PR, Watanabe K, Stringer S, Skene N, Bryois J, Hammerschlag AR, de Leeuw CA, Benjamins JS, Muñoz-Manchado AB, Nagel M, et al. 2019. Genome-wide analysis of insomnia in 1,331,010 individuals identifies new risk loci and functional pathways. Nat Genet 51: 394–403.

Jones SE, Lane JM, Wood AR, van Hees VT, Tyrrell J, Beaumont RN, Jeffries AR, Dashti HS, Hillsdon M, Ruth KS, et al. 2019. Genome-wide association analyses of chronotype in 697,828 individuals provides insights into circadian rhythms. Nat Commun 10: 343.

Juric I, Yu M, Abnousi A, Raviram R, Fang R, Zhao Y, Zhang Y, Qiu Y, Yang Y, Li Y, et al. 2019. MAPS: Model-based analysis of long-range chromatin interactions from PLAC-seq and HiChIP experiments. PLOS Comput Biol 15: e1006982.

Klemm SL, Shipony Z, Greenleaf WJ. 2019. Chromatin accessibility and the regulatory epigenome. Nat Rev Genet 20: 207–220.

Lane JM, Jones SE, Dashti HS, Wood AR, Aragam KG, van Hees VT, Strand LB, Winsvold BS, Wang H, Bowden J, et al. 2019. Biological and clinical insights from genetics of insomnia symptoms. Nat Genet 51: 387–393.

Lane JM, Liang J, Vlasac I, Anderson SG, Bechtold DA, Bowden J, Emsley R, Gill S, Little MA, Luik AI, et al. 2017. Genome-wide association analyses of sleep disturbance traits identify new loci and highlight shared genetics with neuropsychiatric and metabolic traits. Nat Genet 49: 274–281.

Langmead B, Salzberg SL. 2012. Fast gapped-read alignment with Bowtie 2. Nat Methods 9: 357–359.

Li H, Handsaker B, Wysoker A, Fennell T, Ruan J, Homer N, Marth G, Abecasis G, Durbin R, 1000 Genome Project Data Processing Subgroup. 2009. The Sequence Alignment/Map format and SAMtools. Bioinforma Oxf Engl 25: 2078–2079.

Li Q, Brown JB, Huang H, Bickel PJ. 2011. Measuring reproducibility of high-throughput experiments. Ann Appl Stat 5: 1752–1779.

Lindtner S, Catta-Preta R, Tian H, Su-Feher L, Price JD, Dickel DE, Greiner V, Silberberg SN, McKinsey GL, McManus MT, et al. 2019. Genomic Resolution of DLX-Orchestrated Transcriptional Circuits Driving Development of Forebrain GABAergic Neurons. Cell Rep 28: 2048–2063.e8.

Marí n O, Rubenstein JLR. 2001. A long, remarkable journey: Tangential migration in the telencephalon. Nat Rev Neurosci 2: 780–790.

Markenscoff-Papadimitriou E, Whalen S, Przytycki P, Thomas R, Binyameen F, Nowakowski TJ, Kriegstein AR, Sanders SJ, State MW, Pollard KS, et al. 2020. A Chromatin Accessibility Atlas of the Developing Human Telencephalon. Cell 182: 754–769.e18.

Martynoga B, Morrison H, Price DJ, Mason JO. 2005. Foxg1 is required for specification of ventral telencephalon and region-specific regulation of dorsal telencephalic precursor proliferation and apoptosis. Dev Biol 283: 113–127.

Mercader N, Leonardo E, Azpiazu N, Serrano A, Morata G, Martínez-A C, Torres M. 1999. Conserved regulation of proximodistal limb axis development by Meis1/Hth. Nature 402: 425–429.

Nora EP, Lajoie BR, Schulz EG, Giorgetti L, Okamoto I, Servant N, Piolot T, van Berkum NL, Meisig J, Sedat J, et al. 2012. Spatial partitioning of the regulatory landscape of the X-inactivation centre. Nature 485: 381–385.

Nord AS, Blow MJ, Attanasio C, Akiyama JA, Holt A, Hosseini R, Phouanenavong S, Plajzer-Frick I, Shoukry M, Afzal V, et al. 2013. Rapid and Pervasive Changes in Genome-wide Enhancer Usage during Mammalian Development. Cell 155: 1521–1531.

Olsson M, Björklund A, Campbell K. 1998. Early specification of striatal projection neurons and interneuronal subtypes in the lateral and medial ganglionic eminence. Neuroscience 84: 867–876.

Panganiban G, Rubenstein JLR. 2002. Developmental functions of the Distal-less/Dlx homeobox genes. Development 129: 4371–4386.

Pennacchio LA, Ahituv N, Moses AM, Prabhakar S, Nobrega MA, Shoukry M, Minovitsky S, Dubchak I, Holt A, Lewis KD, et al. 2006. In vivo enhancer analysis of human conserved non-coding sequences. Nature 444: 499–502.

Pineault N, Helgason CD, Lawrence HJ, Humphries RK. 2002. Differential expression of Hox, Meis1, and Pbx1 genes in primitive cells throughout murine hematopoietic ontogeny. Exp Hematol 30: 49–57.

Pliner HA, Packer JS, McFaline-Figueroa JL, Cusanovich DA, Daza RM, Aghamirzaie D, Srivatsan S, Qiu X, Jackson D, Minkina A, et al. 2018. Cicero Predicts cis-Regulatory DNA Interactions from Single-Cell Chromatin Accessibility Data. Mol Cell 71: 858–871.e8.

Pollard KS, Hubisz MJ, Rosenbloom KR, Siepel A. 2010. Detection of nonneutral substitution rates on mammalian phylogenies. Genome Res 20: 110–121.

Preissl S, Fang R, Huang H, Zhao Y, Raviram R, Gorkin DU, Zhang Y, Sos BC, Afzal V, Dickel DE, et al. 2018. Single-nucleus analysis of accessible chromatin in developing mouse forebrain reveals cell-type-specific transcriptional regulation. Nat Neurosci 21: 432–439.

Ramírez F, Bhardwaj V, Arrigoni L, Lam KC, Grüning BA, Villaveces J, Habermann B, Akhtar A, Manke T. 2018. High-resolution TADs reveal DNA sequences underlying genome organization in flies. Nat Commun 9: 189.

Reinhardt P, Glatza M, Hemmer K, Tsytsyura Y, Thiel CS, Höing S, Moritz S, Parga JA, Wagner L, Bruder JM, et al. 2013. Derivation and Expansion Using Only Small Molecules of Human Neural Progenitors for Neurodegenerative Disease Modeling. PLoS ONE 8. https://www.ncbi.nlm.nih.gov/pmc/articles/PMC3606479/ (Accessed October 30, 2020).

Ritchie ME, Phipson B, Wu D, Hu Y, Law CW, Shi W, Smyth GK. 2015. limma powers differential expression analyses for RNA-sequencing and microarray studies. Nucleic Acids Res 43: e47–e47.

Robertson KD. 2005. DNA methylation and human disease. Nat Rev Genet 6: 597–610.

Schoenfelder S, Fraser P. 2019. Long-range enhancer–promoter contacts in gene expression control. Nat Rev Genet 20: 437–455.

Schormair B, Zhao C, Bell S, Tilch E, Salminen AV, Pütz B, Dauvilliers Y, Stefani A, Högl B, Poewe W, et al. 2017. Identification of novel risk loci for restless legs syndrome in genome-wide association studies in individuals of European ancestry: a meta-analysis. Lancet Neurol 16: 898–907.

Schulte D, Geerts D. 2019. MEIS transcription factors in development and disease. Dev Camb Engl 146.

Smemo S, Tena JJ, Kim K-H, Gamazon ER, Sakabe NJ, Gómez-Marí n C, Aneas I, Credidio FL, Sobreira DR, Wasserman NF, et al. 2014. Obesity-associated variants within FTO form long-range functional connections with IRX3. Nature 507: 371–375.

Song M, Pebworth M-P, Yang X, Abnousi A, Fan C, Wen J, Rosen JD, Choudhary MNK, Cui X, Jones IR, et al. 2020. Cell-type-specific 3D epigenomes in the developing human cortex. Nature 1–6.

Spieler D, Kaffe M, Knauf F, Bessa J, Tena JJ, Giesert F, Schormair B, Tilch E, Lee H, Horsch M, et al. 2014. Restless legs syndrome-associated intronic common variant in Meis1 alters enhancer function in the developing telencephalon. Genome Res 24: 592–603.

Stadler MB, Murr R, Burger L, Ivanek R, Lienert F, Schöler A, Nimwegen E van, Wirbelauer C, Oakeley EJ, Gaidatzis D, et al. 2011. DNA-binding factors shape the mouse methylome at distal regulatory regions. Nature 480: 490–495.

Stoykova A, Treichel D, Hallonet M, Gruss P. 2000. Pax6 Modulates the Dorsoventral Patterning of the Mammalian Telencephalon. J Neurosci 20: 8042–8050.

Sussel L, Marin O, Kimura S, Rubenstein JL. 1999. Loss of Nkx2.1 homeobox gene function results in a ventral to dorsal molecular respecification within the basal telencephalon: evidence for a transformation of the pallidum into the striatum. Development 126: 3359–3370.

Thakore PI, D’Ippolito AM, Song L, Safi A, Shivakumar NK, Kabadi AM, Reddy TE, Crawford GE, Gersbach CA. 2015. Highly specific epigenome editing by CRISPR-Cas9 repressors for silencing of distal regulatory elements. Nat Methods 12: 1143–1149.

Toresson H, Parmar M, Campbell K. 2000. Expression of Meis and Pbx genes and their protein products in the developing telencephalon: implications for regional differentiation. Mech Dev 94: 183–187.

van de Werken HJG, de Vree PJP, Splinter E, Holwerda SJB, Klous P, de Wit E, de Laat W. 2012a. 4C technology: protocols and data analysis. Methods Enzymol 513: 89–112.

van de Werken HJG, Landan G, Holwerda SJB, Hoichman M, Klous P, Chachik R, Splinter E, Valdes-Quezada C, Öz Y, Bouwman BAM, et al. 2012b. Robust 4C-seq data analysis to screen for regulatory DNA interactions. Nat Methods 9: 969–972.

Walter C, Schuetzmann D, Rosenbauer F, Dugas M. 2014. Basic4Cseq: an R/Bioconductor package for analyzing 4C-seq data. Bioinformatics 30: 3268–3269.

Wickham H. 2016. ggplot2: Elegant Graphics for Data Analysis. Springer-Verlag, New York https://ggplot2.tidyverse.org.

Winkelmann J, Czamara D, Schormair B, Knauf F, Schulte EC, Trenkwalder C, Dauvilliers Y, Polo O, Högl B, Berger K, et al. 2011. Genome-wide association study identifies novel restless legs syndrome susceptibility loci on 2p14 and 16q12.1. PLoS Genet 7: e1002171.

Winkelmann J, Schormair B, Lichtner P, Ripke S, Xiong L, Jalilzadeh S, Fulda S, Pütz B, Eckstein G, Hauk S, et al. 2007. Genome-wide association study of restless legs syndrome identifies common variants in three genomic regions. Nat Genet 39: 1000–1006.

Wonders CP, Anderson SA. 2006. The origin and specification of cortical interneurons. Nat Rev Neurosci 7: 687–696.

Yin Y, Morgunova E, Jolma A, Kaasinen E, Sahu B, Khund-Sayeed S, Das PK, Kivioja T, Dave K, Zhong F, et al. 2017. Impact of cytosine methylation on DNA binding specificities of human transcription factors. Science 356. https://science.sciencemag.org/content/356/6337/eaaj2239 (Accessed June 20, 2021).

Zhang Y, Liu T, Meyer CA, Eeckhoute J, Johnson DS, Bernstein BE, Nusbaum C, Myers RM, Brown M, Li W, et al. 2008. Model-based Analysis of ChIP-Seq (MACS). Genome Biol 9: R137.

Zhao Z, Tavoosidana G, Sjölinder M, Göndör A, Mariano P, Wang S, Kanduri C, Lezcano M, Sandhu KS, Singh U, et al. 2006. Circular chromosome conformation capture (4C) uncovers extensive networks of epigenetically regulated intra-and interchromosomal interactions. Nat Genet 38: 1341–1347.

Ziller MJ, Edri R, Yaffe Y, Donaghey J, Pop R, Mallard W, Issner R, Gifford CA, Goren A, Xing J, et al. 2015. Dissecting neural differentiation regulatory networks through epigenetic footprinting. Nature 518: 355–359.

